# *Chalcolestes color-aeneus* (Odonata, Zygoptera, Lestidae) sp. nov. from Romania

**DOI:** 10.1101/2022.07.27.501700

**Authors:** Anda Felicia Babalean

## Abstract

The paper describes a new Chalcolestes species – *Chalcolestes color-aeneus* collected from Romania. This new species shows morphological similarities with *Chalcolestes viridis* by the metallic iridescent colour of the body and by the yellow to pale brown colour of the lower part of the head; it also shows similarities with *Lestes virens, Lestes sponsa* and *Lestes dryas* by the long fringing hairs on the inferior male annal appendages. *Chalcolestes color-aeneus* can be separated by the other Lestes and Chalcolestes species by the dark coppery colour with green and reddish reflections of the head, of the thorax and dorsal abdomen and especially by the shape and colour of the male anal appendages.

## Introduction

Chalcolestes Kennedy, 1920 is a genus of Zygoptera, consisting of two species: *C. parvidens* (Artobolevskij, 1929) and *C. viridis* (Vander Linden, 1825) (Wildermuth & Martens 2019). Both species are cited in the Romanian fauna – *C. viridis* as *Lestes viridis* (Cîrdei & Bulimar 1965).

## Material and method

One Zygopteran male was accidentally collected into a Barber pitfall trap for Opiliones filled with ethanol 85°, during 05 – 09 August 1998. It is preserved in ethanol 75-80°. The specimen was observed under binocular, both in alcohol and in dry state. For the species identification, the following resources were used:

- Kennedy C. H., 1920 (No. 2), Forty-two hitherto unrecognized genera and subgenera of Zygoptera, The Ohio Journal of Science, vol XXI, No.2: 83-88.
- Cîrdei F., Bulimar F., 1965, Fauna Republicii Populare Române, Insecta, volumul VII, fascicula 5 Odonata, 274 pg.
- Askew R. R., 2004, The dragonflies of Europe (second edition), 308 pg.
- Boudout J-P., Doucet G., Grand D., 2019, Cahier d’identification des libellules de France, Belgique, Luxembourg & Suisse, 151 pg.
- Wildermuth H., Martens A., 2019, Die Libellen Europas, 958 pg.
- Smallshire D., Swash A., 2020, Europe’s Dragonflies – A field guide to the damselflies and dragonflies, pdf-e-resource.

Morphological terminology follows Askew (2004).

## Results

Systematic

Odonata Fabricius 1792

Zygoptera Selys 1840

Lestidae Selys 1840

*Chalcolestes color-aeneus* sp. nov. Babalean (Figures. 1 – 9)

**Fig. 1.**
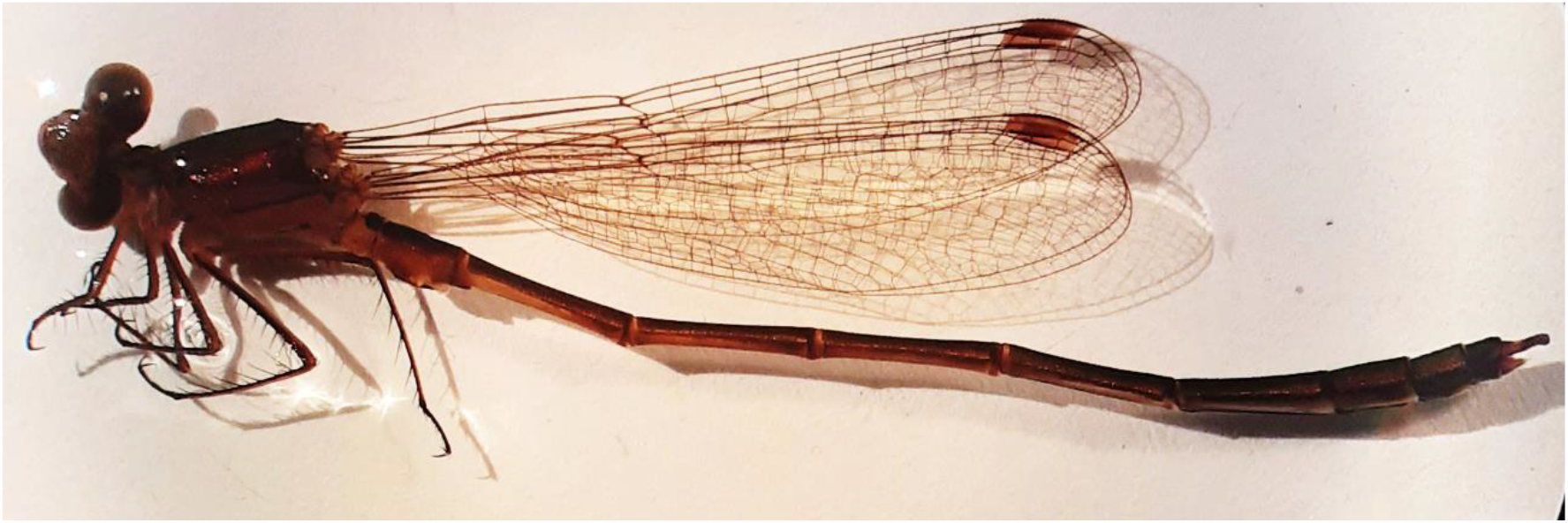
*Chalcolestes color-aeneus* – general aspect in lateral view.

Etymology

The name derives from the colour of the holotype.

Material examined: Holotype male Cce-H-Ro, collected by Anda Felicia Babalean, deposited in author private collection.

### Diagnosis

*Chalcolestes color-aeneus* can be separated by the European Lestes and Chalcolestes species by a set of characters: the dark coppery colour with green and reddish reflections; the uniform colour without spots of the occiput and synthorax; the male anal appendages, especially the shape of the inferior appendages.

### Description of the holotype

*Measurements*: total – 39 mm; head, from the most external part of one eye to the other – 4,46 mm; forewing – 22 mm; hindwing – 21 mm.

*General colour*: dark coppery (bronzy), with green and red metallic iridescence under the light of the binocular both in alcohol and in dry state – Figs. 1, 2, 3, 4.

**Fig. 2.**
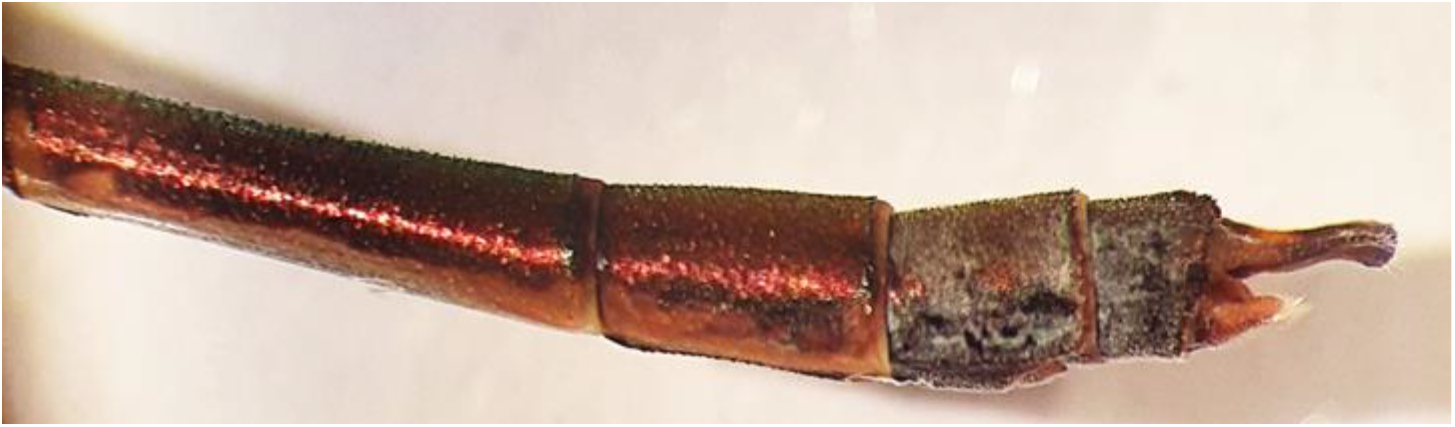
*Chalcolestes color-aeneus* – pruinescence on the last two abdominal segments (in the dry state)

**Fig. 3.**
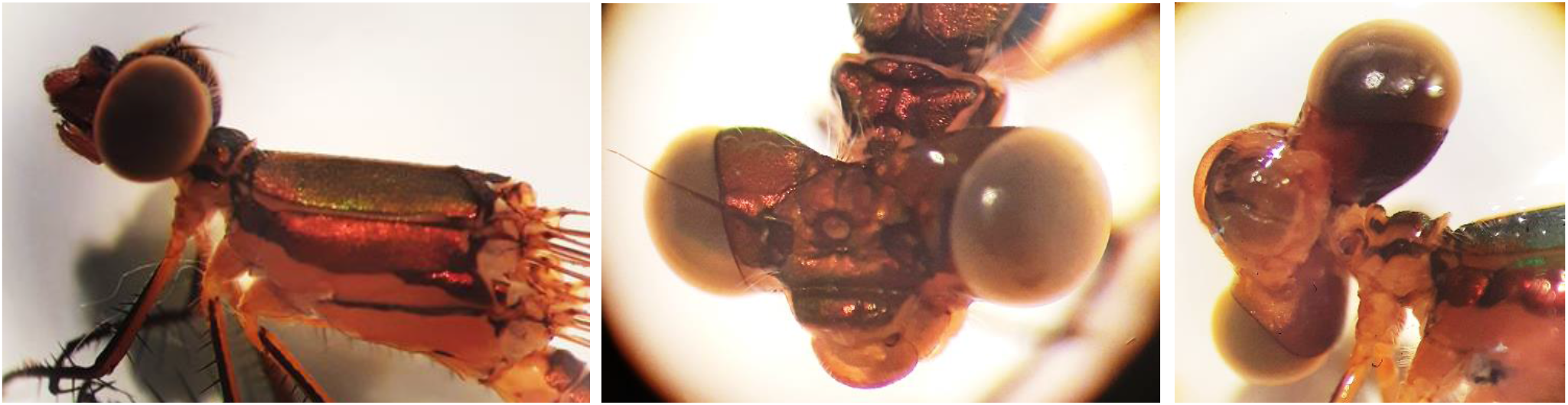
*Chalcolestes color-aeneus* – head and thorax.

**Fig. 4.**
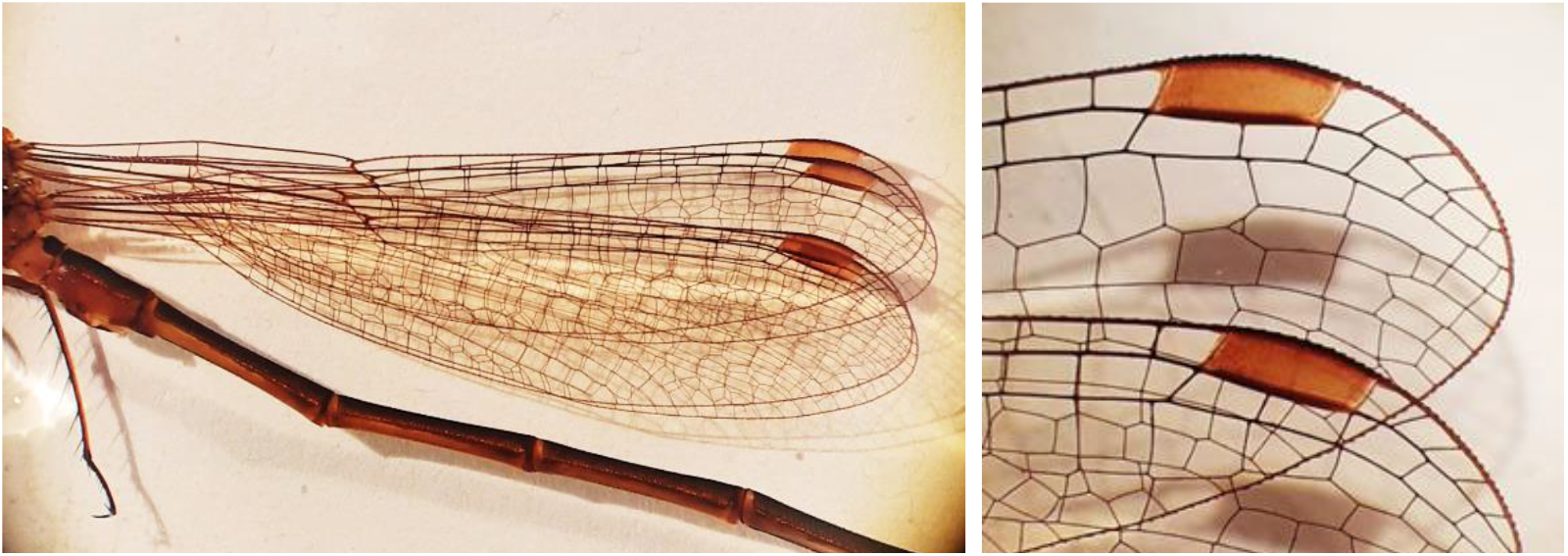
*Chalcolestes color-aeneus* – wings in lateral view, pterostigma.

*Head*: vertex, frons, postclypeus dark bronzy; anteclypeus, labrum, mandible, labium yellowish brown (pale brown in dry state); occiput monocolor, dark bronzy; eyes gray-brown, antenna dark brown – Figs. 1, 3.

*Thorax*: prothorax dark bronzy with yellow spots (yellowish to pale brown in dry state); synthorax dark bronzy, iridescent – Figs. 1, 3.

Head and thorax covered with very thin and long white hairs visible in the dry state.

*Legs*: brown with a yellow stripe; with long spines.

*Wings*: hyaline pale brown, dark-brown veins; pterostigma extending on two cells, brown yellowish with dark upper and lower contours; discoidal cells of the fore- and hindwings almost equal (the difference in size is almost imperceptible); the upper segment of the arculus equals the lower – Figs. 1, 4, 5.

**Fig. 5.**
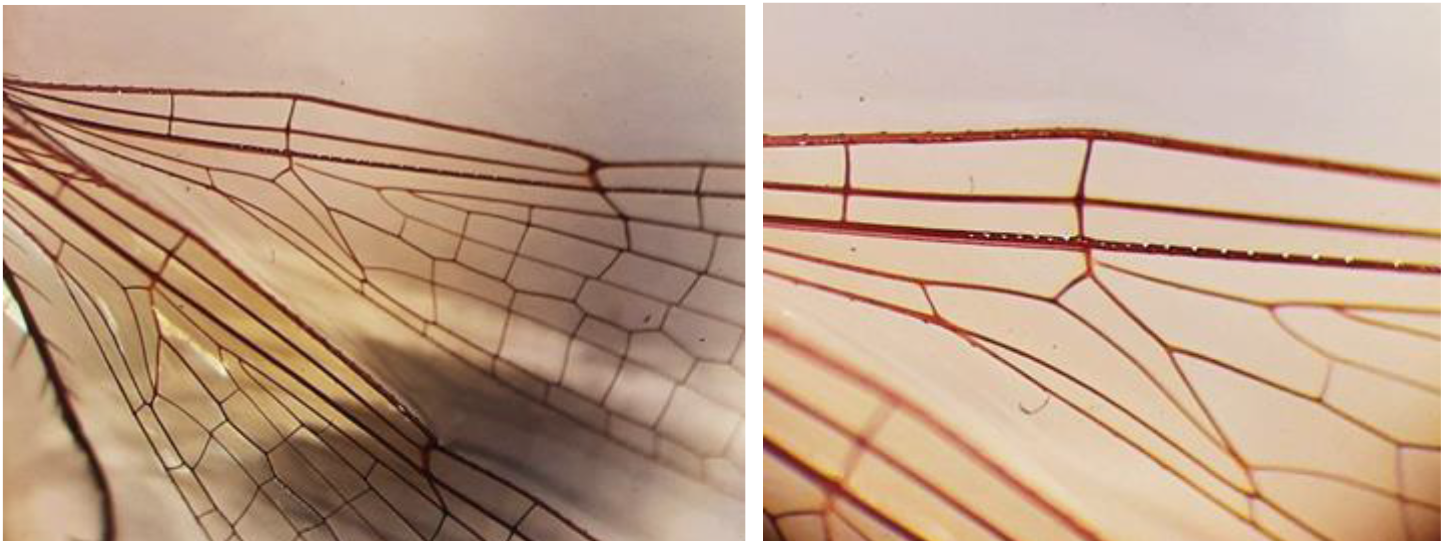
*Chalcolestes color-aeneus* – the discoidal cells and arculus.

*Abdomen*: dark bronzy iridescent on dorsal; yellow in alcohol / pale brown in dry state on ventral, with a large longitudinal brown band on each sternit. Segments S 9 and S10 with a discrete blue pruinescence visible only in the dry state – Fig. 2.

#### Annal appendages

*Superior (external) appendages* – Figs. 6, 7: medially curved, touching each other; blunt tip; the medial (internal) side with a flat denticulated area; external side with small denticles; one internal basal strong spine; colour almost entirely brown with one large yellow basal spot on the external side of each appendage.

**Fig. 6.**
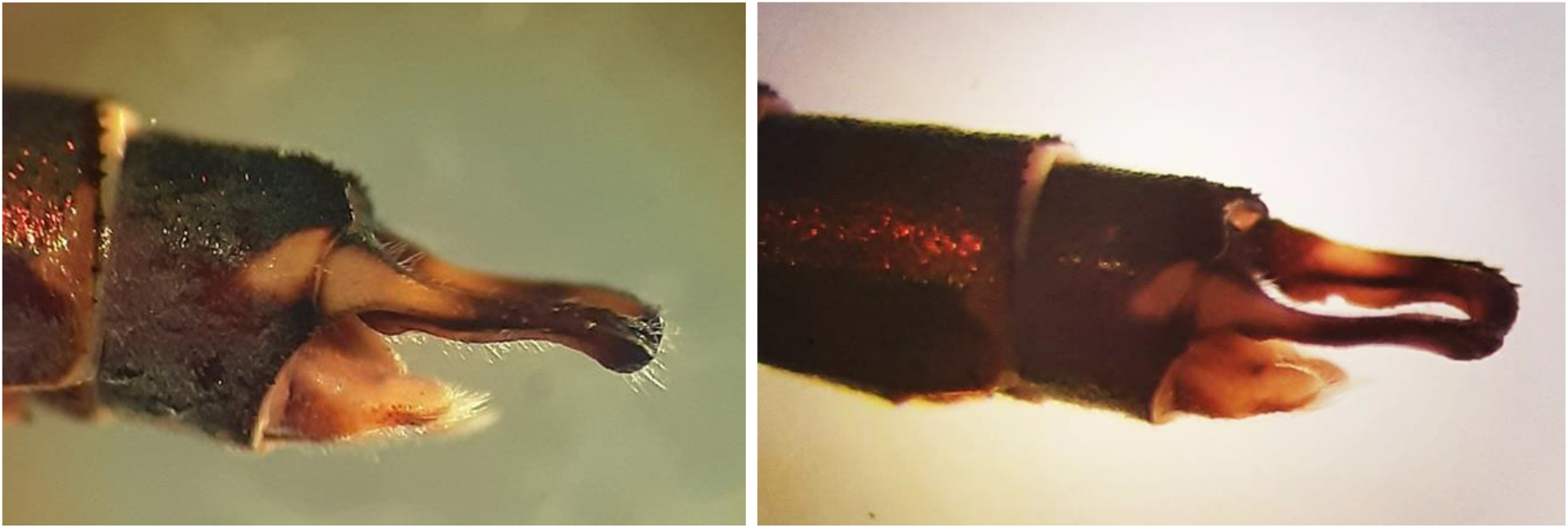
*Chalcolestes color-aeneus* – the anal appendages in lateral view and under different angles.

**Fig. 7.**
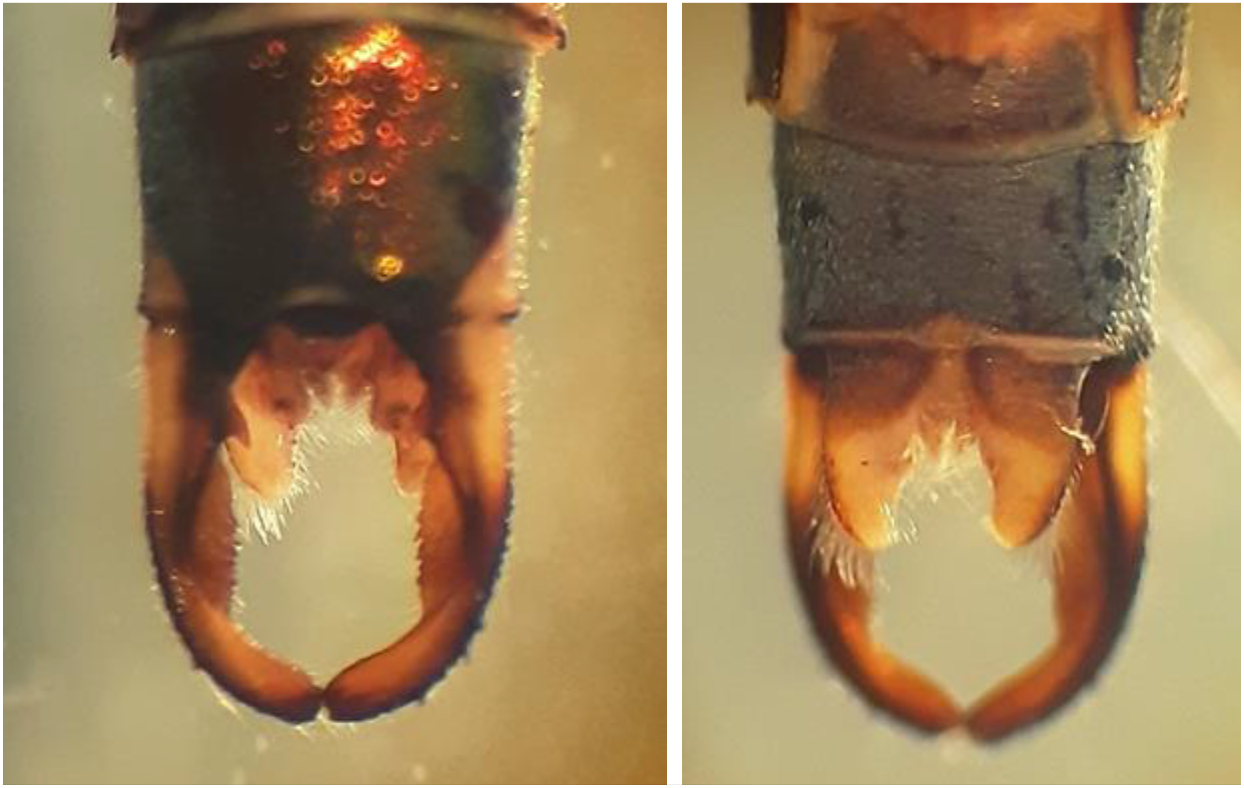
*Chalcolestes color-aeneus* – the anal appendages in dorsal view (left) and in ventral view (right)

*Inferior (internal) appendages* – Figs. 6, 7: lighter colour then the superior appendages; triangular and three-dimensional shape with an internal hairy excrescence (/tubercle/spur); blunt tip; the tip and the external sides covered with long fringing hairs (setae).

*Accessory genitalia* – Figs. 8, 9: hamuli anteriores rounded; vesica spermalis dark brown; the head of the ligula (aedeagus) covered by a membrane and provided with spiniform structures.

**Fig. 8.**
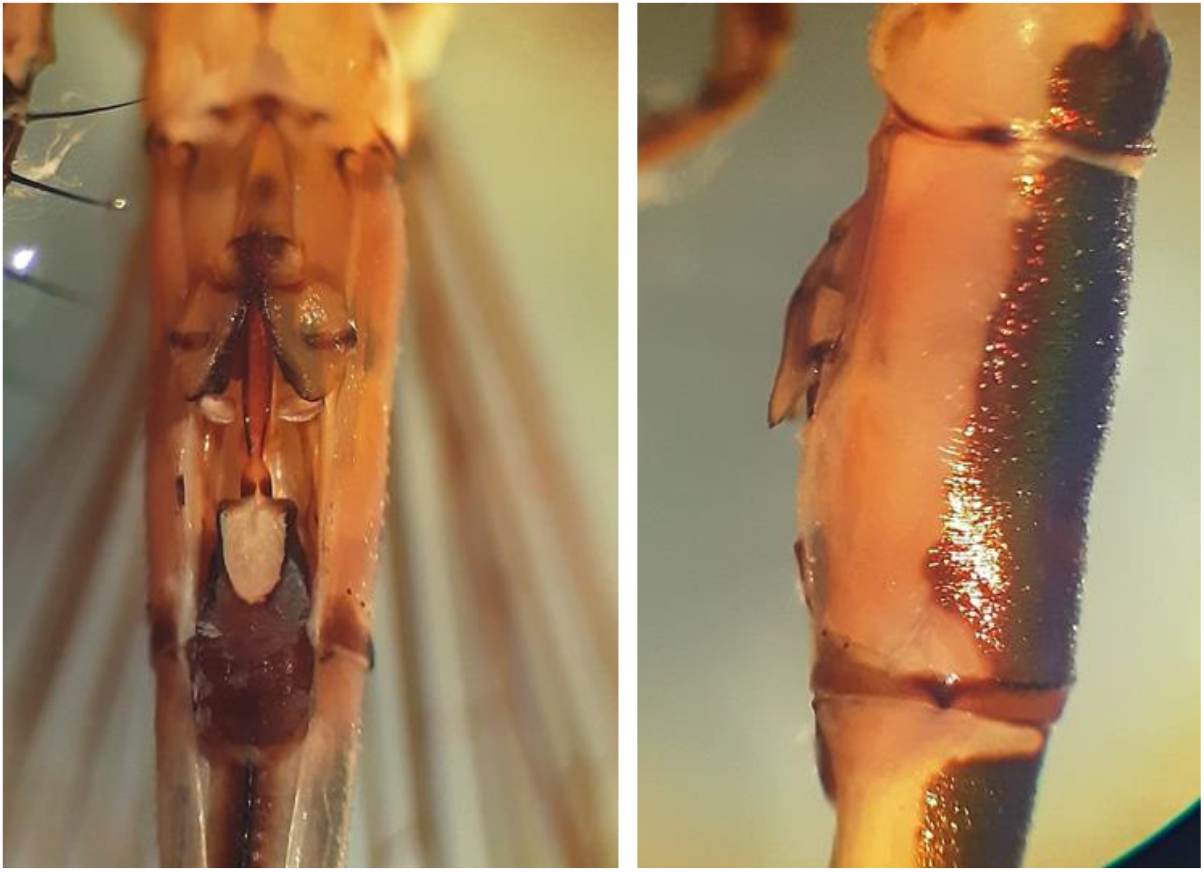
*Chalcolestes color-aeneus* – the accessory genitalia in ventral view (left) and in lateral view (right)

**Fig. 9.**
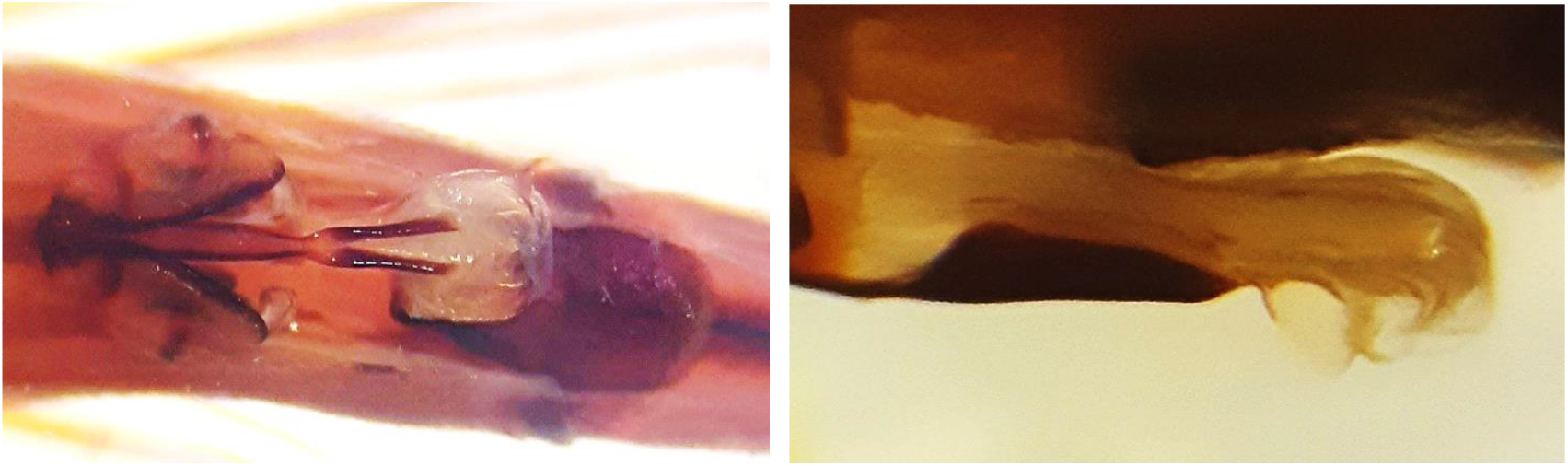
*Chalcolestes color-aeneus* – the aedeagus in ventral view (left) and in lateral view (right)

## Discussions

Compared with other Lestidae (Lestes and Chalcolestes species), *Chalcolestes color-aeneus* shows a high similarity with *Chalcolestes viridis* by the metallic iridescent colour of the body and by the yellowish / pale brown colour of the lower part of the head; it also shows similarities with *Lestes virens*, *Lestes sponsa* and *Lestes dryas* by the long fringing hairs on the inferior male annal appendages (Askew 2004, pg. 59, Figs 38-43). *Chalcolestes color-aeneus* has distinct a set of characters: the dark coppery colour with green and reddish reflections; the uniform colour without spots of the occiput and synthorax; the male anal appendages, especially the shape of the inferior appendages.

The aedeagus is a character used in Odonata systematic and phylogenetic studies, for defining the genera and inferring phylogenetic relationships (for instance Kennedy 1920 No.1). In the most metazoan, the aedeagus (penis, copulatory apparatus) is species specific, being an important mechanism for reproductive isolation of species. In Odonata Zygoptera, it appears that the first mechanism of reproductive isolation of species (to avoid heterospecific breeding) is not the penis, but the “match between male annal appendages and female mesostigmal plates, acting as a mechanical barrier to interspecific tandems” (Cordero-Rivera, Córdoba-Aguilar 2010, citing Robertson & Paterson 1982). The role of the penis is rather for sperm transfer (from male to female) and for sperm removal of the male rivals (Cordero-Rivera, Córdoba-Aguilar 2010). Thus, the reproductive isolation at least of some Odonata species might not be completely, permitting interspecific mating and breeding, with a deep implication in speciation process. Interspecific mating and hybrids appear to be possible and even common (Cordero-Rivera, Córdoba-Aguilar 2010 and included references).

The comparison of the penis head of *Chalcolestes viridis* (Cordero-Rivera, Córdoba-Aguilar 2010, pg. 338, Fig. 15-4(e)) with that of *Chalcolestes color-aeneus* is not possible by simply comparing the photographs.

Recent collecting effort gave no results in finding more specimens males and females of this species. *Chalcolestes color-aeneus* may be an extinct species due to habitat degradation. The odonatological literature reports Zygoptera endemic species restricted to a very small area, sometimes to a single stream, these species being “under imminent threat” (for instance Dijkstra et al. 2007, Vilela et al. 2020).

## Conclusions

Further sampling is important either to complete the description of this new species with the female and larvae or to establish its status as *extinct species.*

